# Multicellular factor analysis of single-cell data for a tissue-centric understanding of disease

**DOI:** 10.1101/2023.02.23.529642

**Authors:** Ricardo O. Ramirez Flores, Jan D. Lanzer, Daniel Dimitrov, Britta Velten, Julio Saez-Rodriguez

## Abstract

Single-cell atlases across conditions are essential in the characterization of human disease. In these complex experimental designs, patient samples are profiled across distinct cell-types and clinical conditions to describe disease processes at the cellular level. However, most of the current analysis tools are limited to pairwise cross-condition comparisons, disregarding the multicellular nature of disease processes and the effects of other biological and technical factors in the variation of gene expression. Here we propose a computational framework for an unsupervised analysis of samples from cross-condition single-cell atlases and for the identification of multicellular programs associated with disease. Our strategy, that repurposes multi-omics factor analysis, incorporates the variation of patient samples across cell-types and enables the joint analysis of multiple patient cohorts, facilitating integration of atlases. We applied our analysis to a collection of acute and chronic human heart failure single-cell datasets and described multicellular processes of cardiac remodeling that were conserved in independent spatial and bulk transcriptomics datasets. In sum, our framework serves as an exploratory tool for unsupervised analysis of cross-condition single-cell atlas and allows for the integration of the measurements of patient cohorts across distinct data modalities, facilitating the generation of comprehensive tissue-centric understanding of disease.

**Graphical Abstract:** 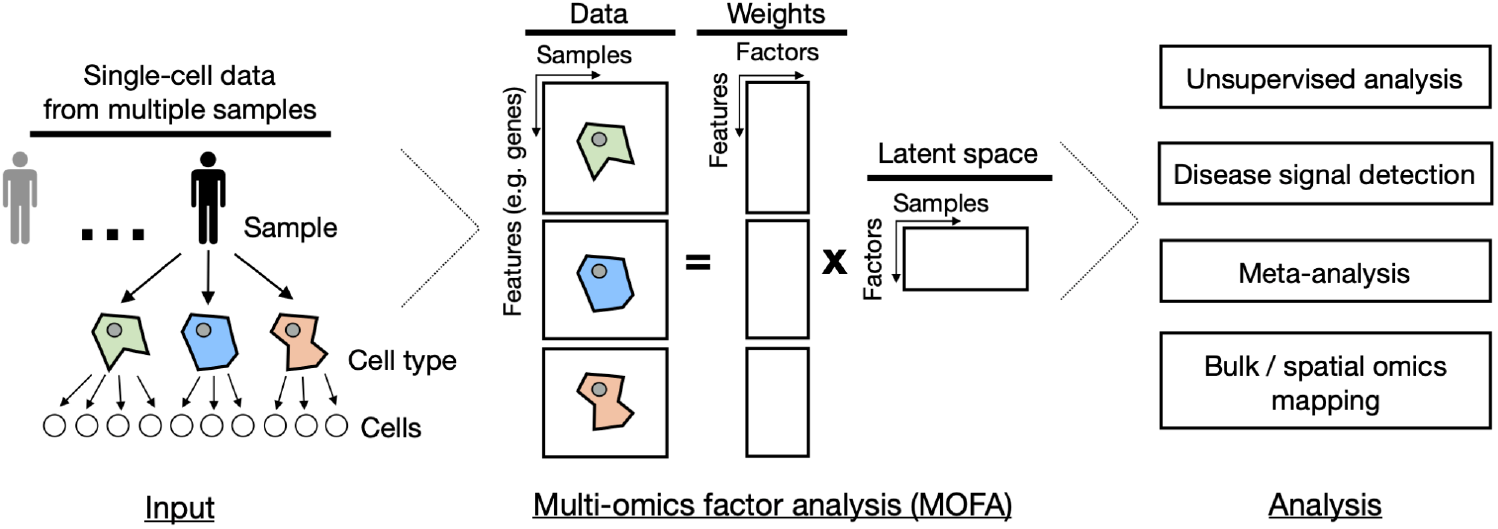

## 1. Introduction

The availability of cross-condition single-cell transcriptomics atlases profiling the pathological state of different tissues and organs in humans has increased during the last years and will continue to expand in different areas of the biomedical field (Rood *et al*, 2022). In these studies a common objective is to compare across groups of samples or conditions the molecular profiles of major cell lineages, commonly referred to as cell-types. Differential gene expression analysis is commonly performed for this task, in which the gene expression of each cell-type is contrasted across various conditions (Crowell *et al*, 2020; Squair *et al*, 2021). This cell-type-centric approach treats each cell-type-specific alteration in disease independently from each other, ignoring coordinated multicellular programs, where particular gene expression changes of one cell-type may relate to the changes of other cell-types. Another limitation of these approaches is that they require a specific definition of cross-condition contrasts *a priori*. Such definitions could disregard other biological and technical variation factors that influence gene expression across cell-types.

A set of novel tissue-centric computational methods have emerged that are helpful in the definition of multicellular programs associated with clinical covariates of interest (Jerby-Arnon & Regev, 2022), and the unsupervised analysis of samples from cross-condition single-cell atlases (Armingol *et al*, 2022; Mitchel *et al*, 2022). These methods are extensions of matrix factorization that aim to reduce the dimensionality of the data while retaining most of the variability. In contrast to classic approaches, such as principal component analysis, these methods are capable of dealing with higher order representations, such as the ones from single-cell data, where a sample is described by a collection of different cell-types. A key element of these methods is that they first transform cross-condition single-cell data into a multi-view representation, in which each view contains the summarized gene expression profile across cells of the same type for each sample.

Although all of these methods can capture coordinated gene expression events across cell-types associated with disease from single-cell data, no current framework has been proposed to map these multicellular programs to other complementary data types such as spatial and bulk omics. Spatial data could be used to understand the spatial regulation of multicellular alterations in disease. Moreover, multicellular programs could be used to deconvolute cell-type-specific gene expression alterations in disease from bulk transcriptomics data, complementing current cell-type deconvolution methods that only estimate cell-type compositions of tissues (Avila Cobos *et al*, 2020). This integrative framework would facilitate the meta-analysis of patient samples across technologies.

Here, we show that multi-omics integration methods, such as Multi-Omics Factor Analysis (MOFA) (Argelaguet *et al*, 2018, 2020), can be repurposed in a straightforward manner to perform similar tissue-centric analyses as the ones performed by the aforementioned methods, since MOFA treats similar multi-view data representations and model objectives to create latent spaces. Moreover, MOFA is a flexible statistical framework that overcomes the limitation of data completeness that some tissue-centric methods enforce (Armingol *et al*, 2022; Mitchel *et al*, 2022), where all samples must contain information in all cell-type views and all cell-type views must contain the same features. In contrast to the aforementioned methods, it also provides the possibility of jointly analyzing independent groups of samples with various classes of cell-type views.

As a case study, we use a collection of acute (Kuppe *et al*, 2022) and chronic human heart failure datasets (Chaffin *et al*, 2022; Reichart *et al*, 2022). We use MOFA for the unsupervised analysis of samples in cross-condition single-cell atlases and the inference of multicellular transcriptional programs associated with technical and biological covariates. We present distinct downstream analyses to relate the inferred multicellular programs to pathway activities and functional cell-states. Moreover, we use spatial transcriptomics to identify the areas in tissues where multicellular disease programs occur. Finally, we use MOFA to meta-analyze single-cell data from multiple patient cohorts to infer multicellular programs that are conserved in independent bulk transcriptomics data. Our analyses represent a flexible multicellular framework that integrates single-cell, spatial, and bulk transcriptomics to analyze cross-condition comparisons to understand tissue alterations during disease. We provide a guided tutorial for the application of multicellular factor analysis to cross-condition single cell atlases in https:/github.com/saezlab/MOFAcell.

## 2. Results

### 2.1 Multicellular factor analysis using MOFA

The generation of a latent space that captures the variability of patients across distinct independent measurements is a task that has been addressed by state-of-the-art multi omics integration methods established for bulk data. The objective of these methods is to integrate independent collections of features (views) measured in the same samples in an unsupervised manner. Hence, we hypothesized that we could repurpose the statistical framework of these multi-view integration methods, such as MOFA (Argelaguet *et al*, 2018, 2020), to describe the variability of samples from single-cell data across cell-types (Figure 1). Based on probabilistic group factor analysis, MOFA can infer a latent space from a collection of cell-type views that contain the summarized gene expression profile of each cell-type per patient (eg. pseudobulk). The variables that form this latent space can be interpreted as coordinated transcriptional changes occurring in multiple cells, here referred to as multicellular programs, providing a tissue-centric understanding of the analyzed sample.

**Figure 1.**
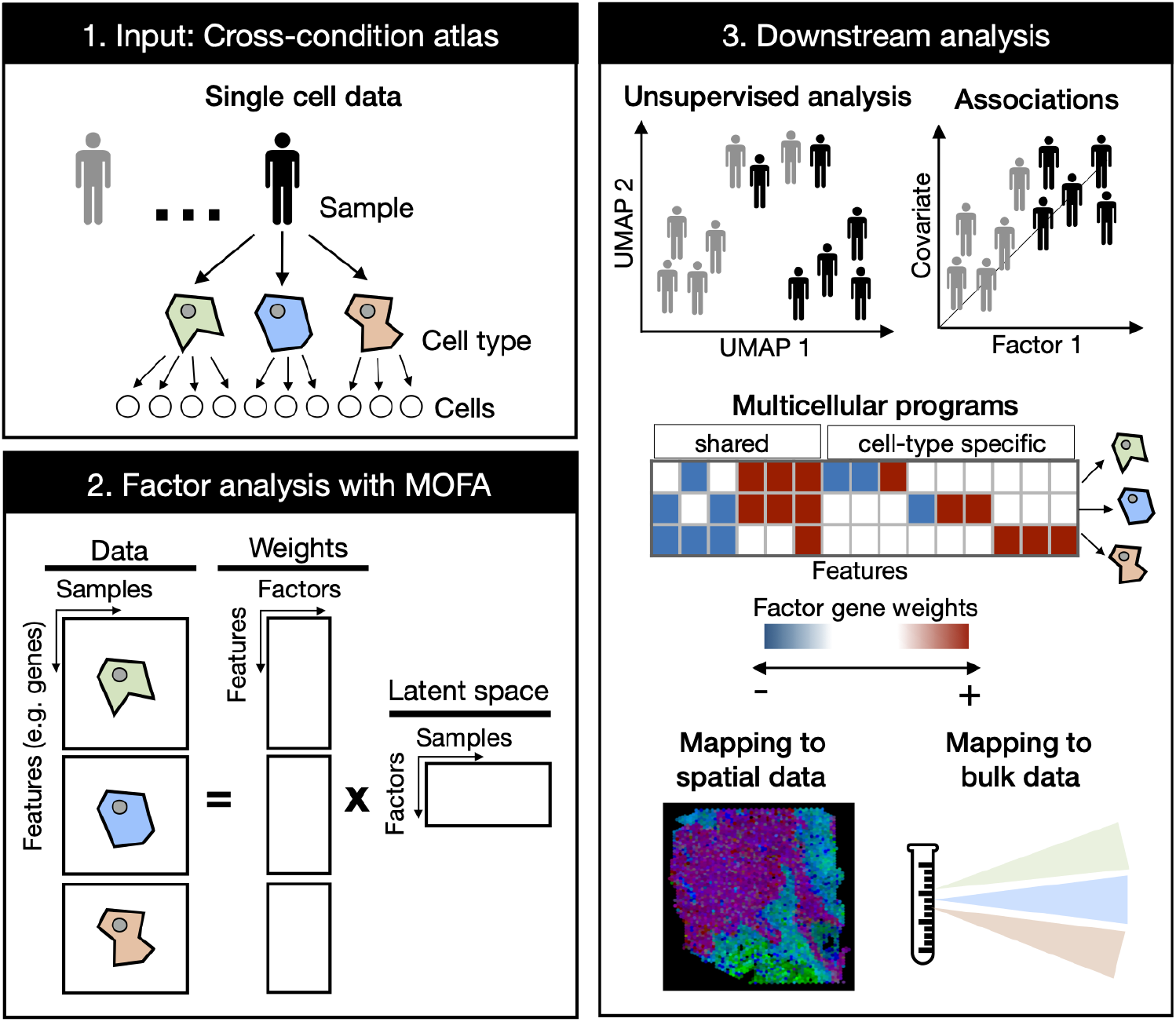
Multicellular factor analysis using MOFA on cross-condition single-cell data. Cross-condition single-cell omics data sample the variability of cells across cell-types, patients, and conditions. The information of these datasets can be summarized as a multi-view representation, i.e. a collection of matrices containing cell-type features across samples. Multicellular factor analysis repurposes multi-omics factor analysis (MOFA) to simultaneously decompose the variability of multiple cell-types and create a latent space that recovers multicellular transcriptional programs. Throughout this manuscript, several applications are presented to show how this analysis can be used for an unsupervised analysis of single-cell data of multiple samples and conditions, for the identification of multicellular disease processes using the inferred latent space, and for a combined analysis of multiple studies across technologies, such as bulk or spatial transcriptomics.

Compared to other methods tailored for the inference of multicellular programs and sample-level unsupervised analysis of single-cell data (Table 1), MOFA allows for a more flexible definition of multi-view integration, since it does not restrict cell-type views to the same features. In addition, MOFA can model samples missing complete cell-type views. MOFA models are computationally efficient (Argelaguet *et al*, 2020) and interpretable, providing measures of the contribution of each view and feature in the construction of the latent space. Finally, building upon these properties, the MOFA-derived cell-type specific gene weights can be used to generate disease signatures that can be mapped to other modalities such as spatial and bulk omics (Figure 1).

**Table 1.** Comparison of methods for multicellular analysis

| Method | Statistical method | Decomposition | Input |  |  | Output |  |  |
| --- | --- | --- | --- | --- | --- | --- | --- | --- |
|  |  |  | Data representation | Data type | Non-overlapping features across views/layers | Multicellular programs | Unsupervised analysis of samples | Evaluation of the quality of the latent space |
| DIALOGUE (Jerby-Arnon & Regev, 2022) | Penalized matrix decomposition followed by multi-level modeling | Linear | Multi-view | Summarized gene expression across cell-types | Yes | Yes | No | No |
| scITD (Mitchel *et al*, 2022) | Tucker decomposition | Linear | Tensor | Summarized gene expression across cell-types | No | Yes | Yes | Yes |
| Tensor cell2cell (Armingol *et al*, 2022) | Tensor Component Analysis | Linear | Tensor | Coexpression of ligand and receptors of pairs of sender and receiving cell-types | No | Yes | Yes | Yes |
| MOFA (Argelaguet *et al*, 2020, 2018) | Probabilistic group factor analysis | Linear | Multi-view | Summarized gene expression across cell-types | Yes | Yes | Yes | Yes |

### 2.2 Multicellular factor analysis for an unsupervised analysis of samples in single-cell cohorts

To show that MOFA could be repurposed to perform an unsupervised multicellular analysis of samples profiled with single-cell or nuclei RNA-seq, we fitted a MOFA model to a cross-condition atlas of human myocardial infarction previously generated by us (Kuppe *et al*, 2022). This atlas profiles distinct phases of myocardial remodeling after infarction, which is a multicellular compensatory process that involves the coordination of multiple cell-types for the maintenance of the heart’s function after injury. After quality control, this dataset contained 27 left-ventricle heart single-nuclei samples of three tissue conditions across seven cell-types previously annotated: myogenic (n = 13), fibrotic (n = 5) and ischemic (n = 9) (Figure 2a). The seven cell-types, previously profiled and annotated across samples, included cardiomyocytes (CMs), fibroblasts (Fib), pericytes (PC), and vascular smooth muscle (vSMCs), endothelial (Endo), myeloid and lymphoid cells. First, we transformed the single-cell data into a multi-view data representation by estimating pseudobulk gene expression profiles for each cell-type across samples (Figure 1b). For each specific cell-type pseudobulk expression matrix, we selected highly variable genes across samples and filtered out lowly expressed and background genes (Methods Section 4.4). We then estimated a shared latent space with six factors (Methods Section 4.5).

**Figure 2.**
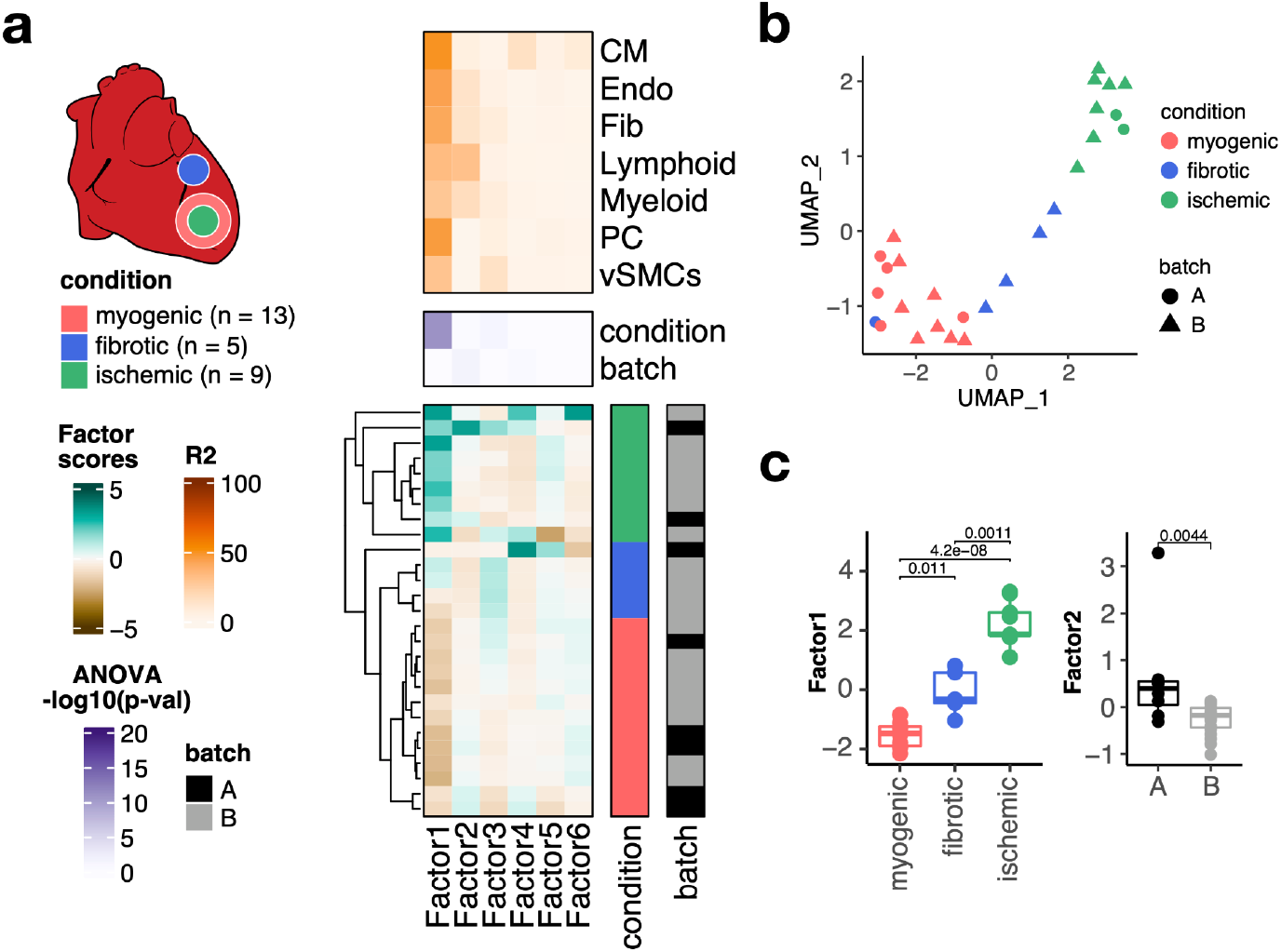
Multicellular factor analysis using MOFA to a single-cell atlas of myocardial infarction. (a) Simplified experimental design of a single-cell atlas of acute heart failure following myocardial infarction from (Kuppe *et al*, 2022). The lower panel shows the factor scores of the 27 samples inferred by the MOFA model. The condition and technical batch label of each sample are indicated next to each row. Samples are sorted based on hierarchical clustering. The middle panel shows the -log10(adj. p-values) of testing for associations between the factor scores and the condition or batch label. The upper panel shows the percentage of explained variance of each cell-type expression matrix recovered by the factor. (b) Uniform Manifold Approximation and Projection (UMAP) embedding of the factor scores from MOFA of each sample in the acute heart failure atlas. (c) Distribution of the scores of Factor 1 across different conditions (left) and of Factor 2 across different technical batches (adj. p-values of t-tests).

The latent space returned by the MOFA model fitted to the single-cell atlas (Figure 2a) explained on average 63.7% of the variability of gene expression of the genes across cell-types. Hierarchical clustering of the samples based on their six factor scores effectively separated ischemic, fibrotic and myogenic-enriched samples. We visualized the sample variability captured by MOFA’s factor scores using an Uniform Manifold Approximation and Projection (UMAP) embedding and multidimensional scaling, and observed similar trends of separation of samples from similar conditions (Figure 2b, Supplementary Figure 1a). From the six recovered factors, Factor 1 was the only factor associated with the previously defined tissue condition labels (ANOVA adj. p-value < 0.05, mean percentage of explained variance across cell-types of 38.8%, Figure 2c), and Factor 2 was associated with the technical label (ANOVA adj. p-value < 0.05, mean percentage of explained variance across cell-types of 11.3%, Figure 2c). Our results suggest that MOFA can be applied to cross-condition single-cell atlases for exploratory unsupervised analysis that allows to detect and prioritize biological signals.

To evaluate the performance of our proposed multicellular factor analysis in the context of related methods, we compared the latent space inferred by MOFA to an analogous one generated with scITD (Mitchel *et al*, 2022) (Supplementary Figure 1b) - to our knowledge, the only other tissue-centric method that provides an interpretable latent space to perform both unsupervised analysis of samples and estimation of multicellular programs (Table 1). First, we observed that compared to MOFA, scITD could only analyze 24 of the 27 samples given the data completeness constraints of their statistical framework based on tensor decomposition. For the shared 24 samples, we evaluated if the latent spaces from both methods could differentiate known labels of patient conditions and technical batches using silhouette scores (Methods Section 4.7). Silhouette scores were comparable across methods for all of the biological and technical labels, except for myogenic-enriched samples which were more similar to each other in MOFA’s latent space (t-test, adj. p-value < 0.01, Supplementary Figure 1c). Since MOFA can handle different sets of genes for each cell-type view, it provides a more flexible framework that enables better control of technical effects in the definition of the latent space, e.g. background genes, compared to methods that enforce data completeness, such as scITD. We quantified the contribution of cell-type specific marker genes, prone to be background, in defining the scITD factor that was associated the most with the patient conditions. We assumed that the definition of the factor would be affected by background noise if TTN, a cardiomyocyte marker gene, and POSTN, a gene expressed in fibroblasts and endothelial cells, would contain high weights across cell-types (Supplementary Figure 1d). As expected, scITD’s absolute gene weights across cell-types were comparable for both marker genes, for example POSTN had a high weight in myeloid cells, a clear background effect since POSTN is not expressed by immune cells. Our results show that the statistical framework of MOFA can be repurposed to infer a latent space from single-cell data that captures the variability of samples across distinct cell-types with comparable performance as the only similar method tailored for this. However, compared to scITD’s framework, MOFA allows for a more flexible definition of cell-type views that better handles missing information and possible technical biases, such as background gene expression.

### 2.3 Multicellular coordinated programs encoded in MOFA’s latent space

To characterize the multicellular molecular processes related to myocardial remodeling captured by MOFA’s inferred latent space from the human myocardial infarction dataset, we inspected and functionally characterized the cell-type specific gene weights that defined Factor 1, the only factor associated with the sample conditions. As previously mentioned, MOFA’s factors can be interpreted as higher-order representations of multicellular programs, i.e. coordinated gene expression changes across cell-types. These patterns encoded in the gene weights of a factor could include gene expression changes shared across multiple cell-types and cell-type specific expression changes (Figure 3a).

**Figure 3.**
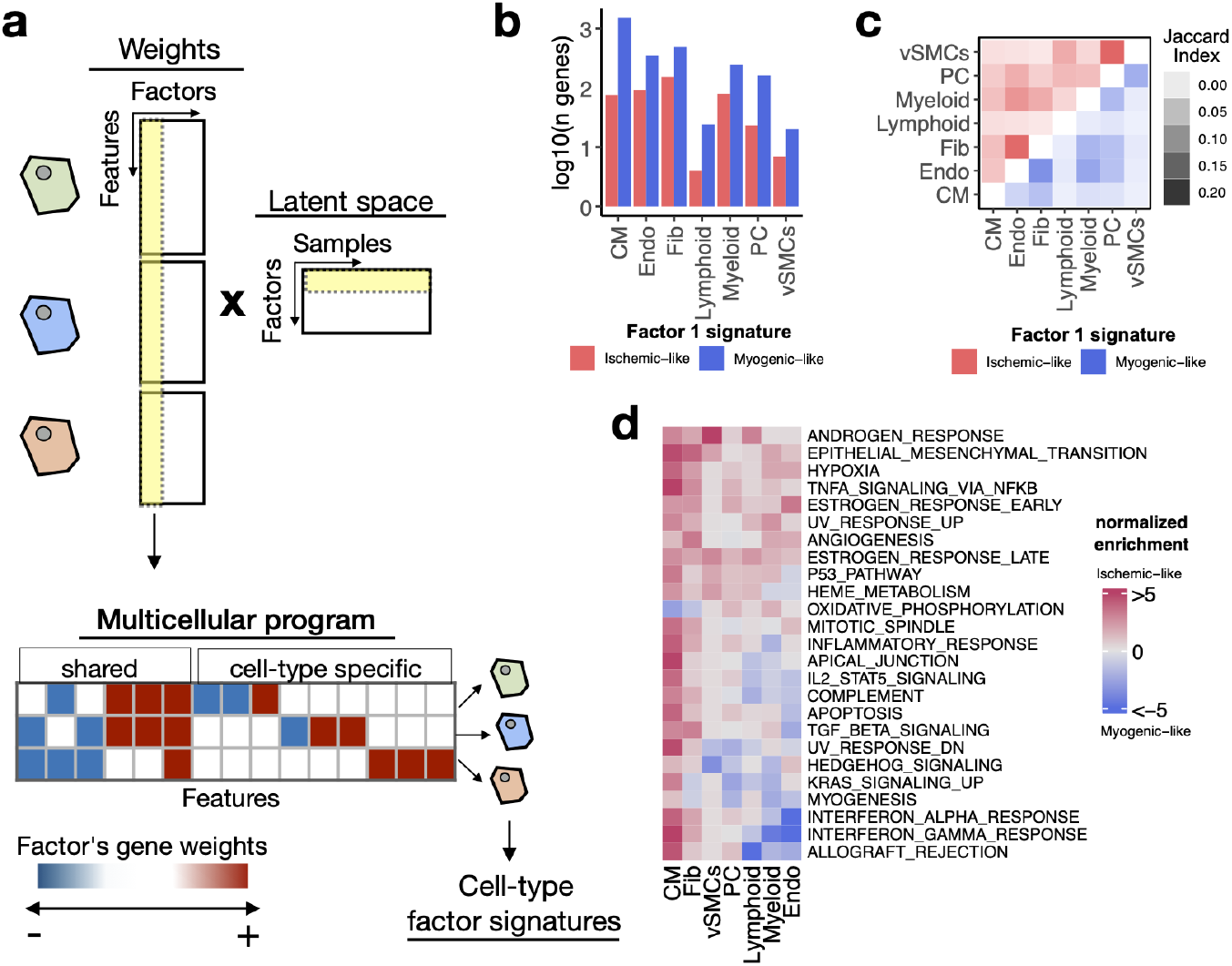
Multicellular programs associated with myocardial remodeling. (a) Each factor forming the latent space reconstructed by MOFA when applied to single-cell data can be interpreted as a higher-level representation of coordinated molecular processes across cell-types, here referred to as multicellular programs. The specific cell-type signatures from these programs can be recovered from the feature weights across cell-types, where the expression changes of one cell-type relate to the other. Moreover, since each multicellular program associates with the variability of samples (eg. differentially active across conditions), cell-type signatures can also be interpreted in the same context. (b) Log10(Number of genes) for each cell-type factor 1 signature associated with the sample conditions of the human myocardial infarction data. Signatures were divided into ischemic-like or myogenic-like signatures based on the weight of each gene. (c) Jaccard Index across myogenic-like (upper triangle) and ischemic-like (lower triangle) cell-type factor signatures associated with the sample conditions of the human myocardial infarction data. (d) Functional enrichment of MSigDB’s hallmarks in cell-type signatures. Enrichment is quantified as normalized weighted means, using the gene weights of each cell-type signature. Top 25 pathways based on mean absolute enrichment score are shown.

First, from the collection of 3,136 unique highly variable genes used in the model across cell-types (Supplementary Figure 2a), we observed that after filtering by importance (Methods section 4.8), the median number of genes associated with Factor 1 per cell-type was 322. Additionally, 12% of the genes associated with Factor 1 were relevant for more than a single-cell-type, suggesting that the multicellular coordinated gene expression associated with myocardial remodeling captured by MOFA is mainly dominated by cell-type-specific processes (Supplementary Figure 2b). To better distinguish between multicellular processes associated with myogenic and ischemic-enriched samples, we simplified the gene weight matrix of Factor 1 into positive and negative cell-type-specific factor gene signatures (Methods section 4.8). Given the positive association of the scores of Factor 1 with ischemic heart samples, the positive and negative cell-type signatures can be understood as ischemic and myogenic signatures, respectively. We observed that cell-type myogenic signatures (median size across cell-types = 243) were larger than the ischemic ones (median size across cell-types = 76), indicating a general trend of downregulation of gene expression of the cells in the myocardium after ischemic injury (Figure 3b). Additionally, we observed little overlap between myogenic and ischemic cell-type factor gene signatures across cell-types (Jaccard index of 0.06 and 0.04, for ischemic and myogenic signatures, respectively, Figure 3c). Altogether, our observations suggest that the multicellular transcriptional alterations upon myocardial infarction captured by the MOFA model represent mostly cell-type specific processes, with a small subset of general processes shared between cell-types.

Functional characterization of the cell-type specific myogenic and ischemic factor gene signatures revealed known cellular processes of cardiac remodeling upon myocardial infarction (Figure 3d). Enrichment of MSigDB’s hallmarks (Liberzon *et al*, 2015) showed that ischemic signatures captured mainly a multicellular response to hypoxia and inflammation across the majority of the cells, together with enrichment of fibrotic processes and angiogenesis. These expected disease processes are associated with tissue damage and cell death upon myocardial infarction which was also captured by the enrichment of the apoptosis pathway in cardiomyocytes. Myogenic signatures mainly were enriched by homeostatic oxidative phosphorylation processes in cardiomyocytes and fibroblasts, together with specific processes of endothelial cells regarding responses to interferons and TGFb activities. Our results suggest that the multicellular programs encoded in MOFA’s factors provide tissue-level descriptions that facilitate the generation of hypotheses related to disease processes, without the need for independent statistical tests per cell-type.

### 2.4 Cell-type-specific factor gene signatures relate to cell-state abundance

We next quantified to what extent the cell-type-specific factor gene signatures recapitulated the emergence of functional cell-states, which are cells of the same lineage but with distinct functional profile (eg. myofibroblasts). To test for an overrepresented signal of cell-states in each cell-type factor signature, we enriched marker genes of cell-states of cardiomyocytes, fibroblasts, endothelial and myeloid cells presented in our previous work (Kuppe *et al*, 2022). We observed across cell-types that myogenic cell-type factor signatures had an overrepresentation of marker genes of cell states that increased in abundance in myogenic samples, compared to ischemic and fibrotic ones (Figure 4a-left, hypergeometric test adj. p-value < 0.05). In contrast, ischemic signatures were enriched by marker genes of cell-states that increased in abundance in ischemic and fibrotic samples (Figure 4a-right, hypergeometric test adj. p-value < 0.05). These results align with the expected effect of pseudobulk profiles, where the gene expression signal of the most abundant cells is prioritized. Overall, we showed that cell-type-specific factor gene signatures captured transcriptional changes related to the change in compositions of functional cell-states as a consequence of the disease context.

**Figure 4.**
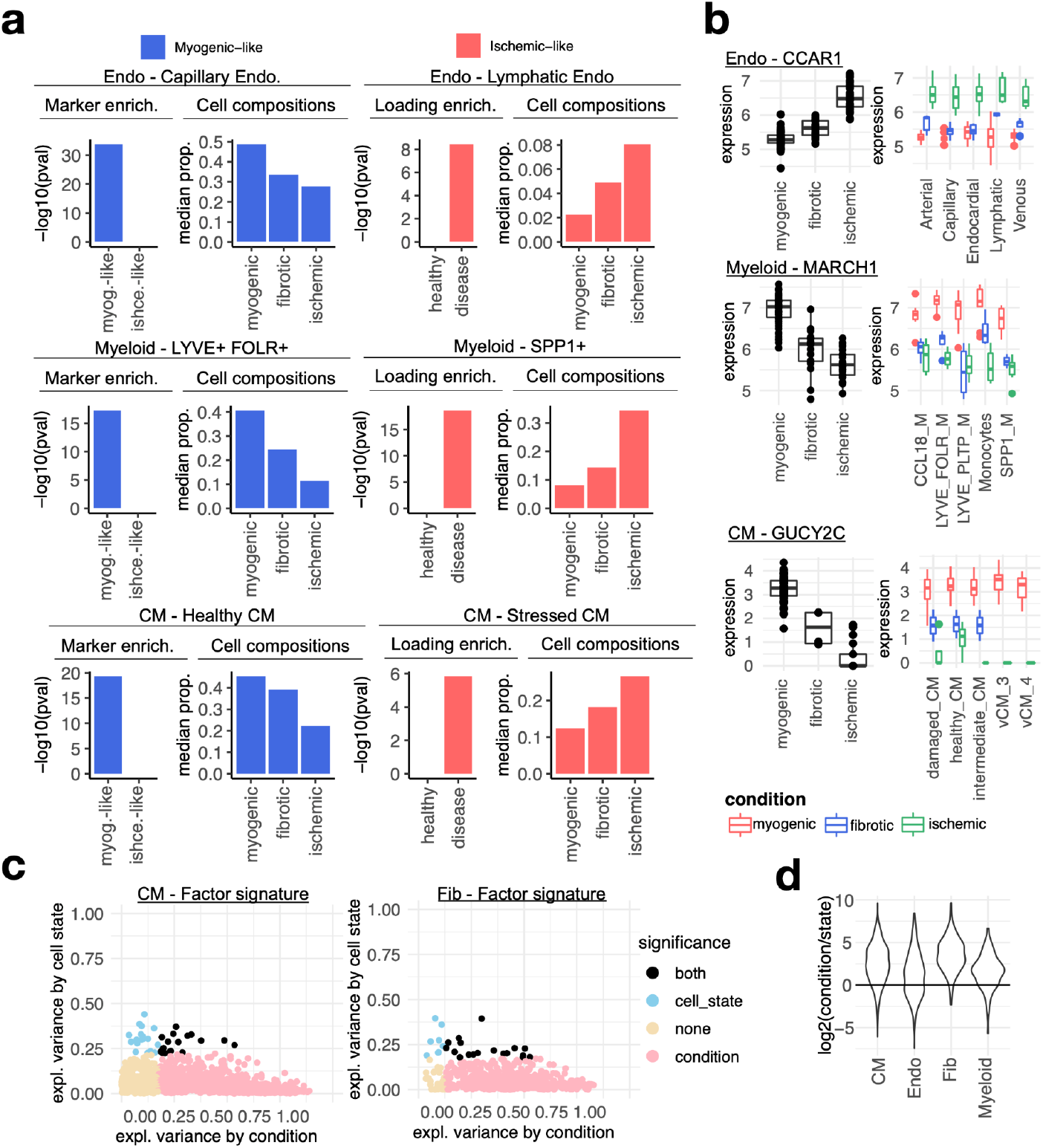
Cell-state dependent and independent transcriptional deregulations upon myocardial infarction. (a) Cell-state markers captured in cell-type factor 1 signatures of endothelial (Endo) and myeloid cells, and cardiomyocytes (CM). In each panel is shown the adj. p-value of the enrichment of cell state markers of the indicated state in cell-type specific factor signatures (left) and the group median proportion of cell-states across patient groups calculated from single-cell data (right). The most overrepresented state in myogenic (left column) and ischemic (right column) samples is shown per cell-type. (b) Distribution of log-normalized gene expression across samples of the human myocardial infarction atlas of CCAR1, MARCH1, and GUCY2C from pseudobulk profiles of Endo and myeloid cells, and CMs, respectively. The left column is the pseudobulk expression of the gene in a cell-type for each sample, while the right column comes from the pseudobulk expression of the gene in a cell-state. (c) Proportion of variance (eta-squared) of gene expression of genes belonging to the factor 1 signatures of cardiomyocytes and fibroblasts (Fib) explained by the patient’s conditions or cell states. Colors represent significance of the association of pseudobulk expression variability to patient’s conditions or cell states (ANOVA adj. p-value < 0.05). (d) Distribution of the log2 ratios across cell-types between the proportion of variance explained by the patient condition and cell-states. Each point in each distribution is the ratio of a gene of the cell-type factor 1 signature (see c).

### 2.5 Cell-type-specific factor gene signatures are dominated by cell-state independent transcriptional changes

Next, we questioned if a global transcriptional response to ischemic injury across cells within a lineage could be recovered from the cell-type-specific factor gene signatures. We hypothesized that while the emergence of cell-states is a valid abstraction of the molecular processes related to disease, there may be transcriptional changes that are independent from cell-states and represent a global alteration of cells within the diseased tissue. This would mean that within a cell-type, the deregulation of a gene as a consequence of a disease context can be traced across cell-states.

We tested this hypothesis by contrasting the proportion of variance of gene expression that could be explained by the samples’ condition and the cell-state classes within each cell-type factor gene signature (Methods section 4.11). Across all cell-type-specific factor gene signatures, we observed differentially expressed genes between conditions (ANOVA adj. p-value < 0.01) that were conserved across cell-states (Figure 4b-c). We observed that in general, across all cell-type signatures, a greater proportion of variance of gene expression was explained by the samples’ condition, rather than the cell-state (one-sample-t-test adj. p-value < 0.01, Figure 4d), suggesting that the genes defining the multicellular latent variable associated with myocardial remodeling recovered by the MOFA model capture both cell-state dependent and independent transcriptional changes. Moreover, these results suggest that while certain cell-states emerge during myocardial infarction, cells within a tissue and lineage partake in a shared global transcriptional response to injury.

### 2.6 Spatial mapping of multicellular coordinated programs

In addition to the functional characterization of multicellular programs with pathway activities and cell-states as presented in the previous sections, complementary data types, such as spatial transcriptomics, can be used to better understand their coordination in intact tissues. Thus, we next mapped the cell-type-specific factor signatures associated with myocardial remodeling to the collection of 28 paired spatial transcriptomics slides (10x Visium) that were generated together with the single nuclei data used in the previous sections (Methods section 4.2, 4.12). Given our previous observation that cells within a lineage could respond to cardiac injury in a cell-state independent manner, we reasoned that the expression of multicellular transcriptional programs associated with myocardial remodeling could be distributed in larger areas in ischemic and fibrotic tissues compared to myogenic-enriched specimens.

For each spatial transcriptomic slide, we calculated the relative area where myogenic and ischemic cell-type factor gene signatures were expressed, using the cell-type composition information in each location. As hypothesized, we observed that across cell-types, except for pericytes and lymphoid cells, the expression of ischemic programs occur in larger areas and with a bigger magnitude in ischemic samples compared to myogenic samples (t-test adj. p-value < 0.05, Figure 5a-b). Similarly, the expression of myogenic programs of fibroblasts and cardiomyocytes was more abundant in myogenic samples compared to the ischemic ones (t-test adj. p-value < 0.05, Figure 5a-b). Compared to myogenic tissues, fibrotic ones expressed ischemic programs of cardiomyocytes and fibroblasts in larger areas, while their myogenic programs were expressed in smaller areas (t-test adj. p-value < 0.05, Figure 5a-b). These results are in line with the expected disease trajectory of myocardial infarction, which progresses from an acute response to injury to chronic compensation that makes the tissues more similar to a healthy myocardium. Our analyses showed that the multicellular program associated with myocardial remodeling captured by MOFA relates to the extent to which myogenic and ischemic cell-type programs are expressed in tissues. Moreover, our mapping strategy provides a complementary analysis strategy for the integration of single-cell and spatial data.

**Figure 5.**
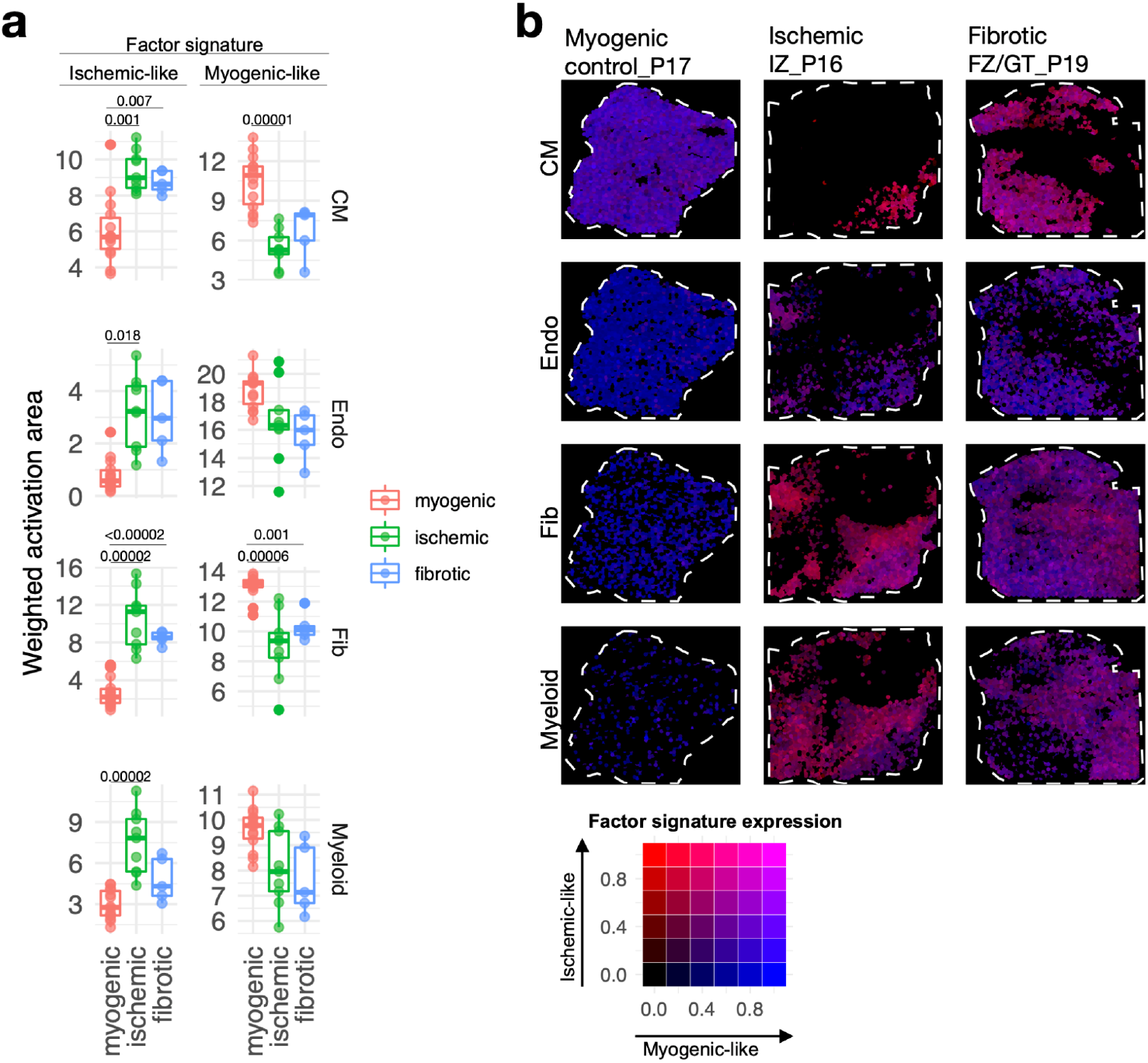
Spatial mapping of multicellular programs associated with myocardial remodeling. (a) Quantification of the relative area of the expression of cell-type specific factor 1 signatures separated by their gene-weights in spatial transcriptomics data of human myocardial infarction for cardiomyocytes (CM), endothelial cells (Endo), fibroblasts (Fib), and myeloid cells. Adjusted p-values of t-tests between conditions are shown for differences where the value was lower than 0.05 (n = 28). (b) Spatial mapping of the activation of CM, Fib, Endo and myeloid cell-type factor 1 signatures in representative examples of myogenic, ischemic and fibrotic samples. The color of each spot represents the combination of the expression of myogenic and ischemic-like programs mapped to the Red-Green-Blue color space (red = ischemic-like, blue = myogenic-like). Patient identifiers as in (Kuppe *et al*, 2022) are indicated in the slide header.

### 2.7 Multicellular factor analysis for the meta analysis of single-cell atlases of heart failure

To show that our proposed framework could be extended to jointly analyze not only multicellular programs but also independent patient cohorts, we performed a meta-analysis of publicly available chronic heart failure single nuclei atlases. Our assumption was that the shared latent space inferred by MOFA would represent multicellular programs that are conserved across distinct etiologies and independent studies. We created multi-view representations of two different single nuclei studies of heart failure across seven cell-types as previously described (Methods section 4.2 and 4.13). The first study (Chaffin2022) encompassed 42 single nuclei cardiac samples profiling healthy myocardium (n = 16) and end-stage heart failure both from dilated (n = 11) and hypertrophic cardiomyopathies (n = 15) (Chaffin *et al*, 2022). The second study (Reichart2022) profiled 79 cardiac samples of healthy myocardium (n = 18) together with samples of dilated (n = 52), non-compaction (n = 1), and arrhythmogenic right ventricular (n = 8) cardiomyopathy (Reichart *et al*, 2022).

After homogenizing the cell-type annotations, we identified shared highly variable genes per cell-type across studies and estimated a joint latent space of six factors using MOFA’s extension for group modeling in addition to study-specific models to define a baseline (Methods section 4.13, Supplementary Figure 4a-c). Baseline study-specific models captured a mean total amount of explained variance across cell-types of 43% for both datasets (Supplementary Figure 4a-b). In contrast to the study-specific models, the joint model had a reduction of 2.8% and 4.1% for Chaffin2022 and Reichart2022, respectively, suggesting that joint-modeling had no critical effects on the construction of the multicellular latent space (Supplementary Figure 4c). We visualized the distribution of samples across studies using the scores of the first two factors of the joint model, which suggested a separation of failing and non-failing hearts regardless of their etiology (Figure 6a). In the study-specific models, we could associate the distinct patient conditions with a mean percentage of explained variance across cell-types of 23% for both studies (ANOVA adj p-value < 0.05, Supplementary Figure 4a-b). The joint model had an increased percentage of 8.6% for Chaffin2022 and 4.2% for Reichart2022, supporting the idea that the inclusion of multiple studies could better define conserved disease signals (Supplementary Figure 4c). From the six factors reconstructed in the joint model, three factors associated with the patient conditions, from which one (factor 1) discriminated failing and non-failing hearts in both studies, and two factors (factor and 3) were only associated to the differences between conditions in Reichart2022’s (t-test adj. p-value<0.05, Figure 6b, Supplementary Figure 4c). Functional characterization of the cell-type-specific factor 1 signatures revealed known multicellular processes active in failing hearts, such as the activation of JAK-STAT, TGFb and WNT signaling pathways, related to inflammatory and fibrotic processes, together with the reactivation of fetal programs (Liew & Dzau, 2004) (Figure 6c). Our results showed that multicellular factor analysis can be applied to samples coming from distinct patient cohorts for an unsupervised meta-analysis of the transcriptional coordinated responses of a tissue in distinct disease contexts of heart failure.

**Figure 6.**
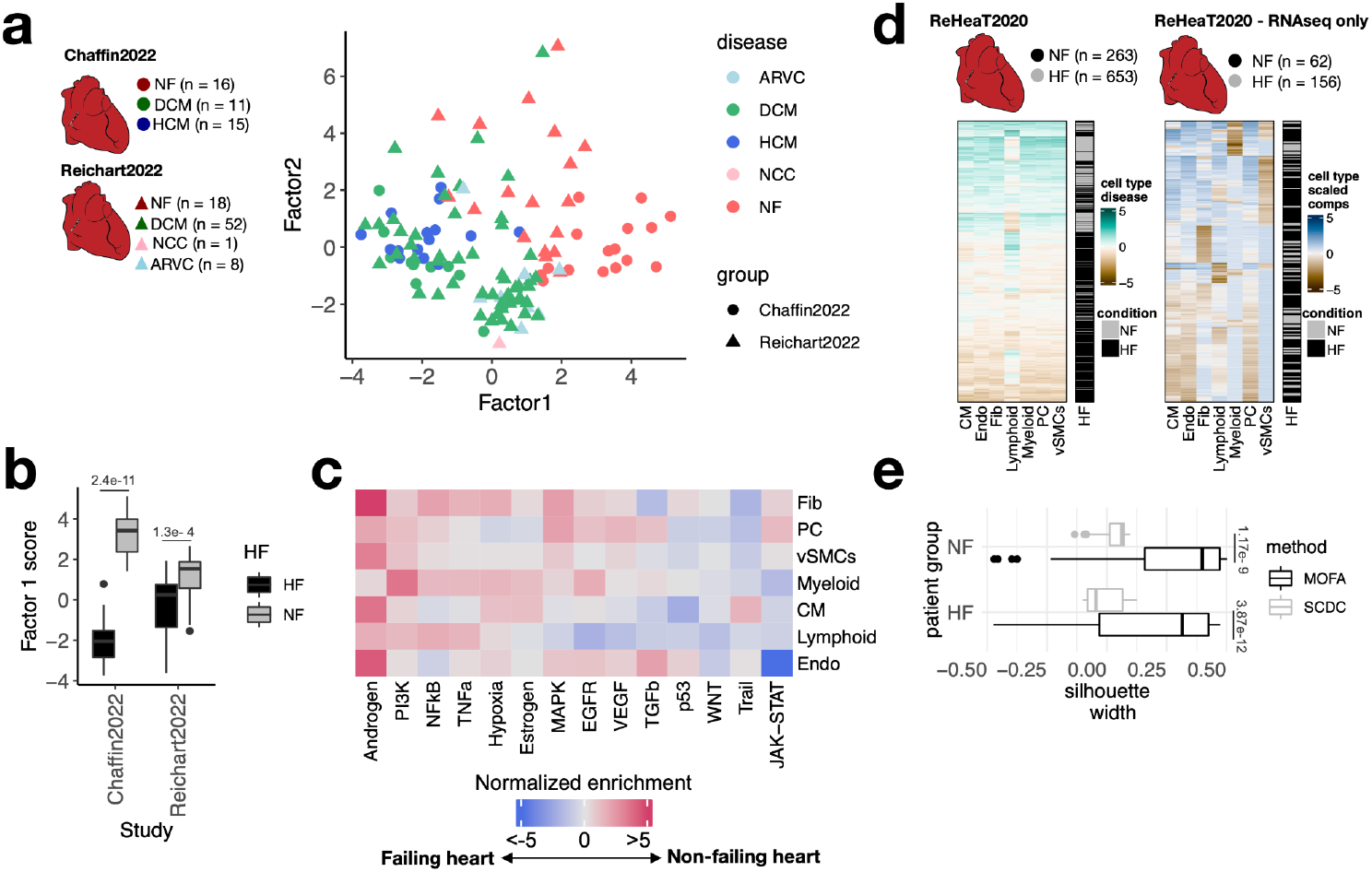
Multicellular factor analysis for the meta-analysis of patient cohorts across technologies. (a) Distribution of patient samples of two single-cell chronic heart failure studies, Chaffin2022 and Reichart2022, based on the first two factors estimated by a grouped multicellular factor analysis model. (b) Distribution of the patient samples of Chaffin2022 and Reichart2022 across the group multicellular factor 1, separated by their heart failure condition: failing (HF) and non-failing (NF) hearts. Adjusted p-values from t-tests are shown. (c) Pathway activities estimated from the gene weight matrices of the group multicellular factor 1 using PROGENy. CM = cardiomyocytes, Fib = fibroblasts, Endo = Endothelial, vSMCs = vascular smooth muscle cells, PC = pericytes. (d) Hierarchical clustering of bulk transcriptomic samples from ReHeaT using scaled cell-type factor 1 signatures (left) or scaled cell-type compositions (center-log-ratio transformed) as estimated by SCDC (right). Heart failure status is indicated for each row: failing (HF) and non-failing (NF). (e) Distribution of silhouette widths of each RNA-seq sample from ReHeaT grouped by their heart failure. Silhouette widths were calculated either by cell-type factor 1 signatures (MOFA) or scaled cell-type compositions (center-log-ratio transformed) (SCDC). Adjusted p-values of t-tests are shown.

### 2.8 Mapping multicellular programs to bulk transcriptomics reveals conserved disease signals across technologies

Finally, we proposed an alternative bulk transcriptomics deconvolution approach of disease signals based on our multicellular factor analysis. We assumed that a bulk expression profile is convoluted by both the cell-type compositions and the coordinated multicellular response of all cell-types of the tissue and that the contribution of each cell-type in the latter effect could be quantified by the enrichment of cell-type specific factor signatures. To test if heart failure multicellular processes could be traced in independent bulk transcriptomics data, we mapped cell-type specific heart failure factor 1 signatures estimated from our previously described joint model of Chaffin2022 and Reichart2022 to an independent collection of 16 bulk heart failure transcriptomics studies (ReHeaT) (Ramirez Flores *et al*, 2021) encompassing 916 human heart samples profiled with microarrays and RNA-Seq (Methods section 4.14). Additionally, from the subset of RNA-seq studies in ReHeaT, we estimated cell-type compositions using established bulk deconvolution methods coupled with the heart human cell atlas as a reference (Litviňuková *et al*, 2020). To justify the selection of the deconvolution method used, we tested the performance of MuSiC (Wang *et al*, 2019), SCDC (Dong *et al*, 2020) and Bisque (Jew *et al*, 2020) in the task of deconvoluting cell compositions from pseudobulk profiles of Chaffin2022 and Reichart2022. We observed that SCDC had the highest median Pearson correlation and the least median root-mean-squared error with the true compositions across studies (median Pearson correlation = 0.84, median root-mean-squared error = 0.108), thus we used this method for the deconvolution of ReHeaT’s studies (Supplementary Figure 4d).

Hierarchical clustering of cell-type specific signature scores across samples in ReHeaT showed a general conservation of the heart failure signature identified from single-cell data, in which bulk failing-hearts had negative signature scores across cell-types (Figure 6d, left). We observed that the conservation of the heart failure signal in bulk samples was independent of their estimated cell compositions, since hierarchical clustering of cell-type compositions led to a greater mixing of failing and non-failing samples (Figure 6d, right), which was quantified using silhouette scores in only RNA-seq samples (t-test, adj. p-value < 0.05, Figure 6e). Given that the datasets in ReHeaT are of various sample sizes, we then tested if cell-type-specific factor signature scores and cell-type compositions separated failing from non-failing hearts in each study individually. In 13 of the 16 bulk studies, we observed a congruent difference in the expression of at least one cell-type-specific factor signature between failing and not failing hearts (t-test adj. p-value < 0.1). Additionally, in 9 of these studies we could differentiate failing and not failing hearts using all of the cell-type-specific factor signatures except for lymphoid cells (t-test adj. p-value < 0.1). In comparison, differential cell-type compositions of at least one cell-type between failing and non-feiling hearts were only observed in 2 of 7 RNA-seq studies (Supplementary Figure 4e). These results show that the multicellular responses associated with heart failure estimated from single-cell data are transferable to different patient cohorts and data modalities. Moreover, the effective mapping of cell-type specific gene programs in bulk samples, suggest that transcriptional profiles from whole tissues are not only driven by highly abundant cell-types, for example cardiomyocytes and fibroblasts in the heart. Our proposed analysis allowed us to meta-analyze over 1000 human heart samples and provides an opportunity to re-analyze bulk data beyond cell-type compositions, serving as a validation ground of single-cell cohorts of smaller size.

## 3. Discussion

Despite the high costs of single-cell technologies, it is expected that in the next few years single-cell datasets encompassing hundreds of patients will be generated. These data hold the promise to allow us to better characterize molecular alterations during disease. Consequently, there is a need for tissue-centric frameworks that on the one hand enable an unsupervised analysis of samples across cell-types and on the other hand provide estimations of coordinated molecular programs that better reflect the multicellular nature of organs.

In this study, we propose to repurpose the statistical framework of multi-omics factor analysis (MOFA) to estimate cross-condition multicellular programs from single-cell transcriptomics data. We demonstrate that the application of MOFA to collections of pseudobulk expression matrices of major cell-types can generate a latent space that captures technical and biological variability of whole tissue specimens independent of cell-type compositional changes. Our proposed framework facilitates the simultaneous identification of different cell-type alterations in disease, reducing the number of independent statistical tests and contrasts. The interpretability of the model allows it to prioritize shared coordinated transcriptional changes between cell-types, without losing the possibility of identifying cell-type specific alterations. Additionally, the reconstruction metrics provided by the model can be used to identify subsets of cell-types that contribute more to specific clinical covariates of samples. MOFA uses Automatic Relevance Determination (Argelaguet *et al*, 2018) to identify the optimal number of factors forming the latent space, which also facilitates its use. We argue that in comparison to novel methods explicitly built for the modeling of multicellular responses (Armingol *et al*, 2022; Mitchel *et al*, 2022; Jerby-Arnon & Regev, 2022), MOFA has two main advantages: 1) Data completeness is not enforced across samples and cell-type views, which allows to better characterize cell-type specific responses and deal with the technical limitations of cell capture and background noise, and 2) joint-modeling of independent studies to generate a shared latent space for samples, which facilitates the integration, comparison and meta-analysis of multiple patient cohorts.

In an application to a collection of public single-cell atlases of acute and chronic heart failure, we found evidence of dominant cell-state independent transcriptional deregulation of cell-types upon myocardial infarction. This may suggest that while certain functional states within a cell-type are more favored in a disease context, most of the cells of a specific type have a shared transcriptional profile in disease tissues. If part of this shared transcriptional profile is interpreted as a signature of the tissue microenvironment that drives cells in tissues towards specific functions, this result may also indicate that a major source of variability across tissues, besides cellular composition, is the degree in which the homeostatic transcriptional balance of the tissue is disturbed. By combining the results of multicellular factor analysis with spatial transcriptomics datasets, we explored this hypothesis and identified larger areas of cell-type-specific transcriptional alterations in diseased tissues. Given these observations on global alterations upon myocardial infarction, we meta-analyzed single-cell samples from two additional studies of healthy and heart failure patients with multiple cardiomyopathies. Here, we found a conserved transcriptional response across cell-types in failing hearts, despite technical and clinical variability between patients. Further, we could find traces of these cell-type alterations in independent bulk data sets. These observations suggest that our approach can estimate cell-type-specific transcriptional changes from bulk data that, together with changes in cell-type compositions, describe tissue pathophysiology. Altogether, these results highlight how MOFA can be used to integrate the measurements of independent single-cell, spatial, and bulk datasets to measure cell-type alterations in disease.

Our work has a number of limitations. Our proposed framework is dependent on the summarization of the nested design of single-cell data studies using pseudobulk profiles per cell-type, which requires the definition of cell-type ontologies before performing a multicellular analysis, an ongoing effort in the single cell community (Osumi-Sutherland *et al*, 2021). In addition, pseudobulk profiles lead to an information loss since the information of multiple cells is aggregated. However, our observations on the conservation of global responses to disease within cell-types across scales suggest evidence that current pseudobulk approaches still provide a meaningful understanding of tissue function. Furthermore, the linear constraints of the inferred latent space by MOFA restrict the type of gene interactions captured by the model. These limitations, however, are shared across current tissue-centric tailored methods. In contrast, models based on generative deep learning (De Donno *et al*, 2022; Boyeau *et al*, 2022) and the Wasserstein metric (Joodaki *et al*, 2022; Chen *et al*, 2020) can take advantage of single-cell measurements to estimate sample-level heterogeneity, but the interpretability of their estimated latent space is limited in comparison to the MOFA models, where features and cell-types can be associated with each factor.

Although this study focused on applying MOFA to understand multicellular responses in tissues, our results also support the application of similar multi-view models to single-cell data, such as MEFISTO (Velten *et al*, 2022) and MuVI (Qoku & Buettner, 2022). MEFISTO, which analyzes complex time-course experimental designs, could be used to explore multicellular coordinated developmental processes. Additionally, MuVI could improve the interpretation of multicellular coordinated processes by incorporating prior knowledge in the inference of the latent space.

In summary, we contributed with a framework that allows the integration of measurements of independent single-cell, spatial, and bulk datasets to contextualize multicellular responses in disease. Additionally, we provided a guided tutorial on how to repurpose MOFA for a multicellular factor analysis in https:/github.com/saezlab/MOFAcell. Our proposed tissue-centric exploratory analysis is scalable and broadly applicable to any single-cell study profiling multiple samples, and it is not limited to transcriptomics measurements or case-control designs.

## 4. Methods

### 4.1 Multicellular factor analysis

We repurposed the statistical framework of multi-omics factor analysis (MOFA) (Argelaguet *et al*, 2020, 2018) to analyze cross-condition single-cell atlases. These atlases profile molecular readouts (e.g. gene expression) of individual cells per sample, following their classification into groups based on lineage (cell-types) or functions (cell states). We assumed that this nested design could be represented as a multi-view dataset of a collection of patients, where each individual view contains the summarized information of all the features of a cell-type per patient (eg. pseudobulk). In this data representation, there can be as many views as cell-types in the original atlas. MOFA is then used to estimate a latent space that captures the variability of patients across the distinct cell-types. The estimated factors composing the latent space can be interpreted as a collection of multicellular programs that capture coordinated expression patterns of distinct cell-types. The cell-type specific gene expression patterns can be retrieved from the factor loadings, where each gene of each cell-type would contain a weight that contributes to the factor score. Similarly, as in the application of MOFA to multi-omics data, the factors can be used for an unsupervised analysis of samples or can be associated with biological or technical covariates of the original samples. Additionally, the reconstruction errors per view and factor can be used to prioritize cell-types associated with covariates of interest.

### 4.2 Datasets

We applied a multicellular factor analysis to three independent published single-cell atlases of acute and chronic heart failure. To ensure the comparability of the analysis across atlases, we defined a heart cell ontology that included the following cell-types: Cardiomyocytes (CM), fibroblasts (Fibs), endothelial cells (Endos), pericytes (PCs), vascular smooth muscle cells (vSMCs), myeloid and lymphoid cells.

#### Human Myocardial Infarction

Single nuclei RNAseq (sn-RNAseq) gene count expression matrices from 27 human heart tissue samples (patient-area) from our previous work (Kuppe *et al*, 2022) were used. The data was downloaded from the Human Cell Atlas (https:/data.humancellatlas.org/explore/projects/e9f36305-d857-44a3-93f0-df4e6007dc97) and imported into a *SummarizedExperiment v1*.*24*.*0* R object. We used the provided cell-type annotations. Data from adipocytes, neuronal, and proliferating cells were excluded since they were present in fewer than 26 patients. Samples were previously annotated as myogenic-enriched, ischemic-enriched, and fibrotic-enriched, summarizing the distinct physiopathological zones and time-points after human myocardial infarction.

For validation of the relevance of the multicellular factor analysis applied to this dataset, we used the matching 28 spatial transcriptomics slides (10x Visium) provided in the publication. Log-normalized data was generated with *normalize_total* and *log1p* functions from *scanpy v1*.*9*.*1 (Wolf et al, 2018)*. Cell-type deconvolution scores per location, previously computed by cell2location (Kleshchevnikov *et al*, 2022), were used as provided in the Human Cell Atlas entry previously mentioned.

#### Human heart failure caused by dilated and hypertrophic cardiomyopathies (Chaffin2022)

Gene count expression matrices from 42 sn-RNAseq left ventricle cardiac samples profiling healthy myocardium (n = 16) and end-stage heart failure both from dilated (n = 11) and hypertrophic cardiomyopathies (n = 15) were obtained from (Chaffin *et al*, 2022). Data was downloaded from https:/singlecell.broadinstitute.org/single_cell/study/SCP1303/single-nuclei-profiling-of-human-dilated-and-hypertrophic-cardiomyopathy. Cell-type annotations were aligned to our proposed cell-type ontology using regular expressions. Unannotated cells were discarded.

#### Human heart failure caused by dilated and arrhythmogenic cardiomyopathies (Reichart2022)

sn-RNAseq gene count matrices from 79 cardiac samples of healthy myocardium (n = 18), together with samples of dilated (n = 52), non-compaction (n = 1), and arrhythmogenic right ventricular (n = 8) cardiomyopathy were collected from (Reichart *et al*, 2022). Left-ventricle data of single nuclei samples were selected from the cellxgene entry: https:/cellxgene.cziscience.com/collections/e75342a8-0f3b-4ec5-8ee1-245a23e0f7cb/private. Cell-type annotations from the authors were adapted to our ontologies using regular expressions and unannotated cells were discarded. Ensembl IDs used in the count matrix were transformed into gene symbols using bioMart v2.50.3 (Durinck *et al*, 2009) and duplicated entries were summed together.

### 4.3 Creation of pseudobulk expression profiles for multicellular factor analysis

Pseudobulk expression profiles were generated for each major cell-type of each independent sample collected in every atlas by summing up the UMI counts of all cells belonging to each of the seven cell-types defined in our ontology. Pseudobulk profiles generated with less than 25 cells were discarded. Genes with less than a minimum of 100 counts in a single sample or detected in less than 25% of the samples were discarded. Data was normalized using the trimmed-mean of M values (TMM) method in edgeR (Robinson *et al*, 2010) with a scale factor of 1 million and log-transformed. Within each atlas, for each cell-type expression matrix, we selected highly variable genes with two strategies. Highly variable genes across samples in the human myocardial infarction atlas were selected for each cell-type using scITD’s adaptation of PAGODA2’s method (norm_variances > 1.5) (Mitchel *et al*, 2022). This was done to enable the comparison between MOFA and scITD. In both of the chronic heart failure atlases, we identified highly variable genes per cell-type using *scran’s v1*.*22*.*1* (Lun *et al*, 2016) *modelGeneVar* function with a biological variance threshold of 0.

### 4.4 Exclusion of background genes from pseudobulk profiles

To avoid including genes belonging to background counts of cell-free mRNA in MOFA’s cell-type views, we limited the genes that could be considered highly variable within each cell-type. For each cell-type, we filtered out all highly variable genes that could be used as markers for any other cell-type. Marker genes of the cell-type from which background genes were filtered out were not considered in the procedure. This filtering procedure reduces the chances of including highly expressed cell-type marker genes that would be more likely to be part of the background counts of all the pseudobulk expression profiles.

In all datasets we identified cell-type marker genes from the differential expression analysis of cell-type pseudobulk expression profiles using *edgeR* (Robinson *et al*, 2010). Genes with a false discovery rate < 0.01 and a log fold change greater than 1 were considered marker genes. Each cell-type was compared against the rest in the model design.

### 4.5 Definition of MOFA models for individual single-cell atlases

A MOFA model with 6 factors was fitted to the collection of pseudobulk cell-type expression matrices, where each cell-type represented an independent view. Gaussian likelihoods were used for each view. Feature-wise sparsity was not forced in the model to obtain the greatest number of genes per cell-type associated with each factor. View data was centered before fitting the model. The six factors were recommended while fitting the model using *MOFA2 v1*.*4*.*0 run_mofa* function to the human myocardial infarction data. MOFA uses Automatic Relevance Determination (Argelaguet *et al*, 2018) to identify the optimal number of factors forming the latent space. For consistency we kept the same number of factors for the rest of the models.

### 4.6 Association of multicellular factor scores to covariates

For a given MOFA model, we associated the factor scores of each patient sample to reported biological and technical covariates in the dataset using analysis of variance (ANOVA). P-values were corrected using the Benjamini-Hochberg procedure. In the human infarction atlas, the patient group and the technical batch label were tested for association with factor scores. In the chronic heart failure atlases, the patient’s condition, etiology, genotype and sex were tested for association with factor scores.

### 4.7 Fitting a scITD model to human myocardial infarction data

To evaluate the capacity of MOFA to fit a multicellular factor analysis, we compared the latent space inferred from the human myocardial infarction dataset to the one recovered by scITD (Mitchel *et al*, 2022), a related method. To fit a scITD model, first, pseudobulk profiles for each cell-type across patients were generated as previously described. Compared to MOFA, scITD represents distinct cell-type views in a tensor, enforcing each cell-type view to contain the same features. Thus, the union of all highly variable genes across cell-types were used in each tensor layer. We used the identical collection of highly variable genes per cell-type previously selected for the MOFA model. Then, a Tucker decomposition (Tucker, 1966) of the pseudobulk tensor was performed with *scITD’s v1*.*0*.*2 run_tucker_ica* function that discarded patient samples with incomplete profiles. A latent space of six factors was recovered to keep consistency with MOFA’s model. Tests for association of inferred factors with clinical covariates were performed with ANOVAs as previously described.

To compare the multicellular latent spaces inferred by MOFA and scITD, we evaluated their ability to differentiate pre-defined histological patient groups and technical batches reported in the human myocardial infarction data. Silhouette scores of each sample were calculated relative to their reported histological class (myogenic, fibrotic or ischemic) and technical batch from an euclidean distance matrix calculated from either the MOFA or scITD factor scores. We performed t-tests to compare the silhouette scores for each histological and technical batch class. P-values were corrected using the Benjamini-Hochberg procedure.

### 4.8 Definition of cell-type specific factor signatures from the MOFA models

After identifying the multicellular factor that associated the most with a covariate of interest (e.g. differences across sample conditions), we defined two factor gene signatures that capture the positive and negative trends of the factor for each cell-type. Given the linear nature of the factor analysis implemented in MOFA, for a cell-type of interest, it is possible to separate the genes with a weight different from 0 into two main classes, positive (> 0) and negative genes (< 0). The interpretation of these two gene sets depend on the direction of association between the factor of interest and the samples’ covariates. For example, if a factor *X* is associated positively with a disease condition, then all genes with a positive weight in a cell-type are also associated positively with the disease condition. All cell-type-specific loadings with absolute values less than 0.1 were set to 0 before the definition of factor signatures.

### 4.9 Functional interpretation of cell-type specific loadings or factor signatures

To functionally characterize the gene loading matrix associated with a factor of interest, we proposed two alternative enrichment analyses based on the mean expression of MSigDB’s hallmarks gene sets (Liberzon *et al*, 2015) and pathway activity footprints (top 500 genes) from PROGENy (Schubert *et al*, 2018). We used *decoupleR’s v2*.*0*.*1 wmean* function (Badia-I-Mompel *et al*, 2022) to calculate weighted mean scores from factor gene weight matrices to have enrichment scores for each cell-type. In the case of PROGENy we used the gene footprints as weights, while for MSigDB’s hallmarks we used unweighted means. The normalized scores of each value were calculated with 1000 permutations.

### 4.10 Estimation of cell-state dependent gene expression changes upon myocardial infarction captured by the multicellular factor analysis

Given a multicellular program explaining the variance of samples represented by the cell-type specific gene loadings of a factor, we quantified to what extent it was associated with the emergence of functional cell-states of the major lineages analyzed. Our hypothesis was that since each cell-type view summarizes gene expression in the form of pseudobulk profiles per sample, then the genes with the highest cell-type specific absolute weights could be associated with cell-states that emerged or increased in a group of samples. We tested this hypothesis in the myocardial infarction dataset by enriching cell-state markers to cell-type-specific factor signatures using hypergeometric tests. Positive and negative signatures were analyzed independently. P-values were adjusted with the Benjamini-Hochberg procedure. For these analyses we only included states defined for cardiomyocytes (CMs), fibroblasts (Fibs), endothelial cells (Endos), and myeloid cells as provided in (Kuppe *et al*, 2022). To calculate cell-state specific markers, within each cell-type, we performed a t-test using *scanpy’s v1*.*9*.*1 rank_genes_groups* function (Wolf *et al*, 2018) at the single-cell level, contrasting the profiles of all cells belonging to one cell-state with the rest of the cells of that cell-type. A gene was considered a marker of cell-state if the log fold change was greater or equal than 0.5 and the adjusted p-value less than 0.05.

### 4.11 Estimation of cell-state independent gene expression changes upon myocardial infarction captured by the multicellular factor analysis

To quantify the extent to which the multicellular programs captured patient variability that was cell-state independent, we assessed whether the expression of a gene, part of a cell-type specific factor signature, was better explained by sample or cell state variability. We hypothesized that within a cell-type-specific factor signature, it would be possible to find genes with uniform gene expression across distinct functional cell-states, which show distinct patterns of expression across distinct groups of samples. This would suggest a general transcriptional shift across cell-states. We tested this hypothesis in the myocardial infarction dataset by performing, within each cell-type, independent ANOVAs to the expression of each gene belonging to its factor signature. The grouping variable was either the patient condition (myogenic, fibrotic, or ischemic) or the cell-state classification. For the former, the ANOVAs were fitted to pseudobulk expression profiles of samples as previously described. For the latter, they were fitted to pseudobulk expression profiles of cell-states across samples within each major lineage (cardiomyocytes, fibroblasts, endothelial and myeloid cells). Profiles generated with less than 25 cells were excluded in both types of tests. Eta-squared values of the grouping variable per gene were used to quantify the amount of variance explained by cell-states or patient conditions. Significance was considered for Benjamini-Hochberg corrected p-values below 0.01. For each gene within each cell-type-specific factor signature, we calculated a log2 ratio between the variance explained by the patient condition and the cell-state as a measure of cell-state independence. For this measure, values over 0 represent a greater explained variance associated to the condition rather than the cell-state. We performed one-sample t-tests on the distributions of the log ratios of explained variance of each gene for each cell-type, to test for general cell-state independence across the factor gene signature.

### 4.12 Spatial mapping of cell-type-specific factor signatures

To map cell-type specific factor signatures to independent spatial transcriptomics (ST) data from the myocardial infarction dataset, we calculated weighted means of gene expression in each location across all ST slides for the positive and negative signatures separately. Normalized weighted mean scores were calculated with *decoupler-py’s v1*.*1*.*0 run_wmean* function (Badia-I-Mompel *et al*, 2022) using as weights the gene loadings of each cell-type specific signature with 100 permutations. Spatial mapping of cell-type factor signatures were only performed in locations where the proportion of the cell-type mapped was equal or greater than 0.1.

To estimate the relative areas across cell-types and ST slides where cell-type-specific factor signatures were expressed, we first assumed that the effective area of a signature for a specific cell-type was defined by the number of spots where the cell-type was present within a ST slide. We consider a cell-type to be present in a location if its proportion was equal or greater than 0.1. Then, for each cell-type-specific factor signature, we counted in how many spots its normalized weighted mean score was greater than two, representing the number of standard deviations from the mean of the distribution of scores from random gene sets. Finally, the relative area of activation of a cell-type-specific factor signature within a slide was calculated as the ratio between the spots with active programs and the effective area.

To visualize the interplay of positive and negative cell-type-specific factor signatures in the ST slides, we encoded the expression of each signature in the red-green-blue (RGB) color space. In this color space, brighter and darker colors represent a high and low expression of a signature respectively and the color combination differentiates different events of co-activation of signatures. To transform the normalized weighted mean estimates into a scale ranging from 0 to 1 so as to be mapped to the RGB space, each cell-type-specific factor signature was normalized by its maximum value across all slides.

### 4.13 Multicellular factor analysis for the integration of independent cohorts

To generate a multicellular factor analysis that integrates the information of independent patient cohorts, we used MOFA’s extension that enables the joint modeling of multiple groups using an extended group-wise prior hierarchy (Argelaguet *et al*, 2020). The main assumption is that the recovered latent space of this group-based analysis will identify factors that explained shared patient variability across studies together with study specific variability. We fitted a grouped MOFA model to two independent chronic end-stage heart failure single-cell studies to identify multicellular programs that differentiated failing and non-failing heart samples. For each study, we generated pseudobulk normalized expression profiles of cell-types for each sample, identified for each cell-type highly variable genes across samples, and filtered out background genes as previously described. Then we selected the collection of highly variable genes per cell-type that were shared across studies and used those to create a joint multi-view representation of both datasets. Finally, we fitted a MOFA model as previously described, but with an additional group variable per sample describing the study of origin and an additional feature-level scaling procedure per study.

### 4.14 Mapping cell-type specific factor signatures to bulk transcriptomics data

To estimate the expression of cell-type specific factor signatures in bulk transcriptomics samples, we estimated normalized weighted mean scores per cell-type signature. For a given bulk transcriptomics study, we calculated normalized weighted mean scores for each cell-type signature using *decoupleR’s v2*.*0*.*1 wmean* function using as weights their gene loadings with 100 permutations. Before the estimation, gene expression data was centered and scaled across samples.

We calculated the expression of the seven cell-type-specific factor signatures associated with heart failure from the joint MOFA model of Chaffin2022 and Reichart2022 for all samples in the 16 RNA-seq and microarray heart failure bulk transcriptomic studies collected in ReHeaT (Ramirez Flores *et al*, 2021). To test for the difference of means of cell-type-specific factor signatures between heart failure and non-failing patients within each study we used t-tests. P-values were corrected using the Benajmini-Hochberg procedure. Significance was assigned to corrected values lower or equal to 0.1.

### 4.15 Benchmarking bulk transcriptomics cell-type deconvolution methods in heart datasets

To evaluate the performance of bulk transcriptomics deconvolution methods in the estimation of cell-type proportions from human heart expression profiles, we benchmarked three methods using the chronic heart failure single-cell datasets. Our benchmark consisted in evaluating the precision of the estimation of cell-type compositions of the samples in Chaffin2022 and Reichart2022 using MuSiC (Wang *et al*, 2019), SCDC (Dong *et al*, 2020) and Bisque (Jew *et al*, 2020) coupled to a healthy heart single-cell reference (Litviňuková *et al*, 2020). First, we selected all apex and left-ventricle snRNA-seq samples from the reference study. Then we manually unified cell type labels and kept all cells belonging to the seven cell-types used in the MOFA models. Next, for both chronic heart failure single-cell datasets, we created pseudobulk profiles of each patient sample summing up gene counts across all cells from all cell types. As a ground truth, we recorded the real proportions of cell-types that were merged into these profiles. We kept the reference and target gene expression matrices in a linear scale and normalized the data using transcripts per million (TPM), as recommended (Avila Cobos *et al*, 2020). Finally, we deconvoluted each pseudobulk sample using MuSiC, SCDC, and Bisque and calculated the Pearson correlation and the root-mean-square error between the estimated and the ground truth cell-type proportions as evaluation metrics.

### 4.16 Cell-type deconvolution of heart failure transcriptomic data sets

Following the results of the benchmark of cell-type deconvolution methods, we estimated the cell-type proportions of CMs, Fibs, Endos, PCs, vSMCs and myeloid cells of all samples across seven RNA-seq heart failure bulk transcriptomic studies collected in ReHeaT (Ramirez Flores *et al*, 2021) using SCDC. Each study was TPM normalized and deconvoluted separately. We used t-tests to test for the difference of means of cell-type compositions between heart failure and non-failing patients within each study. Compositions were transformed to centered-log-ratios using the *clr* function from the *compositions v2*.*0-4* package (van den Boogaart & Tolosana-Delgado, 2008). P-values were corrected using the Benajmini-Hochberg procedure. Significance was assigned to corrected p-values lower or equal to 0.1.

### 4.17 Comparison of cell-type specific factor signatures with cell-type proportions for the separation of failing and non-failing hearts from bulk transcriptomics

To evaluate the biological relevance of mapping cell-type-specific factor signatures to bulk transcriptomics, we compared if these signatures were better at distinguishing failing from non-failing hearts than cell-type proportions, estimated from deconvolution methods applied to bulk transcriptomics data. We assumed that the gene expression profile of a bulk transcriptomics sample could be decomposed by the sample’s cell-type composition and the disease state of each cell-type, as estimated from deconvolution methods and the expression of cell-type-specific factor signatures, respectively. Silhouette scores of each sample across the seven RNA-seq heart failure bulk transcriptomic studies collected in ReHeaT (Ramirez Flores *et al*, 2021) were calculated relative to their condition (failing and non-failing) from an euclidean distance matrix calculated either from their cell-type factor signatures or from their estimated cell-type compositions. Compositions were transformed to centered-log-ratios as previously mentioned before calculating the sample distance matrix. Cell-type factor signatures were scaled across all samples from all studies before calculating the sample distance matrix. We performed t-tests to compare the silhouette scores for each patient condition. P-values were corrected using the Benjamini-Hochberg procedure.

### 4.18 Code and data availability

All scripts related to this manuscript can be consulted here: https:/github.com/saezlab/MOFAcell. A Zenodo entry containing data associated to this manuscript can be accessed here: https:/zenodo.org/record/7660312#.Y_YMLezMIeZ

## 5. Acknowledgements

R.O.R.F. and J.S.R. acknowledge the support of DFG through the CRC 1550 “Molecular Circuits of Heart Disease”. J.D.L. and J.S.R. are supported by Informatics for Life funded by the Klaus Tschira Foundation. D.D. and J.S.R. are supported in part by the European Union’s Horizon 2020 research and innovation program (860329 Marie-Curie ITN “STRATEGY-CKD”). We thank Sebastian Lobentanzer and Olga Ivanova for reading an initial draft of the work and Pau Badia i Mompel, Charlotte Boys, and Robin Fallegger for feedback on the structure of the manuscript. We thank Jovan Tanevski and Ricard Argelaguet for helpful discussions.

## 6. Conflict of interests

JSR reports funding from GSK, Pfizer and Sanofi and fees from Travere Therapeutics, and Astex.

## 7. Authors contributions

R.O.R.F. conceived the project. J.S.R. supervised the project. R.O.R.F. implemented most of the analyses with input from J.D.L, D.D., and B.V. R.O.R.F. generated the figures, with input from all authors. J.D.L. implemented the benchmark of bulk deconvolution methods and conceived the idea of cell-state dependence and independence. All authors interpreted the results. R.O.R.F. wrote the manuscript with input from all authors.

## Supplementary Materials

### Supplementary Files

**Supplementary Figure 1.**
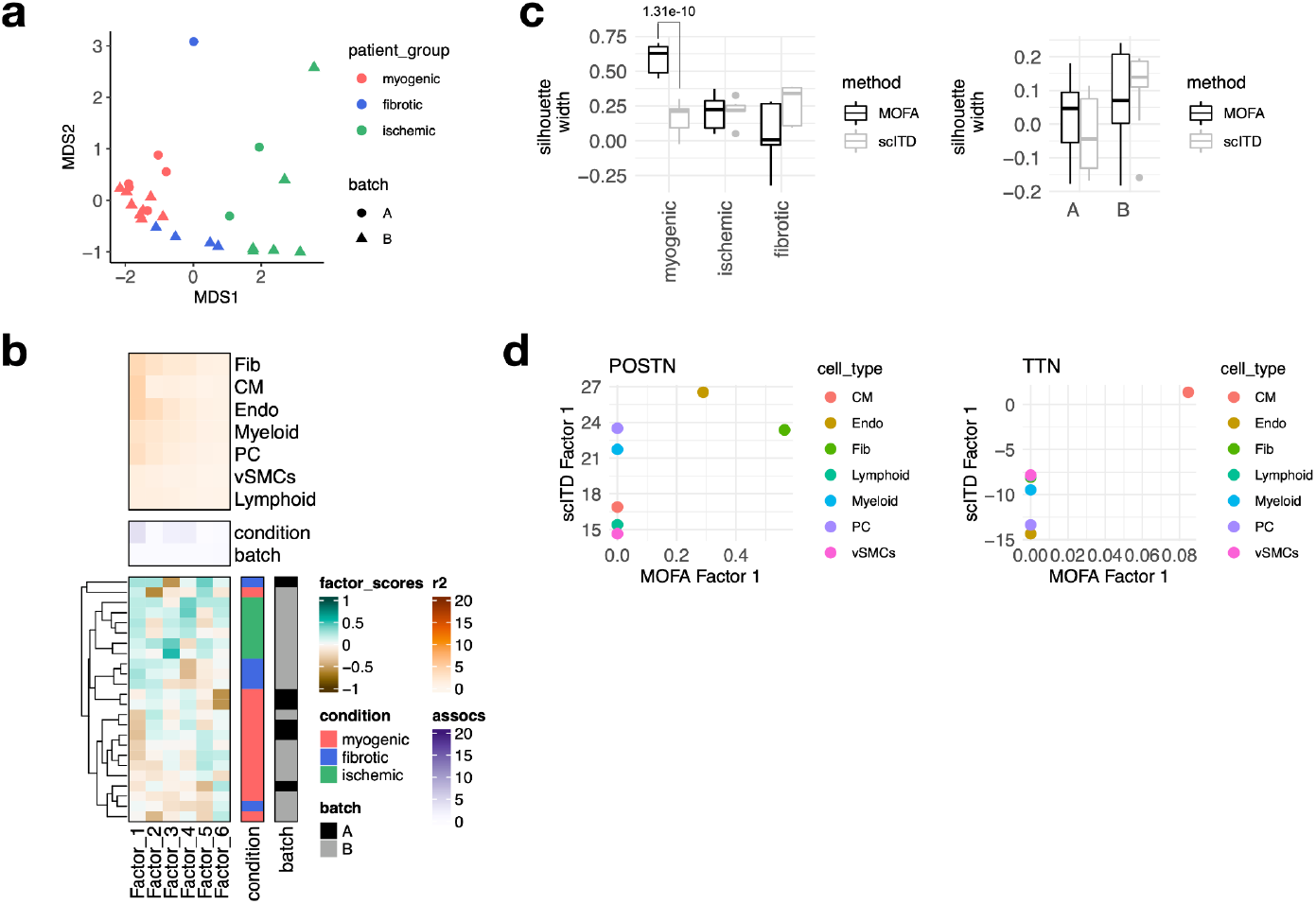
Estimation of a multicellular latent space of acute heart failure using MOFA and scITD. a) Multidimensional scaling (MDS) embedding of the factor scores from MOFA of each sample in the acute heart failure atlas. b) scITD Tucker decomposition of the acute heart failure atlas. The lower panel shows the factor scores of the 24 samples inferred by the scITD model. The condition and technical batch labels of each sample are indicated next to each row. Samples are sorted based on hierarchical clustering. The middle panel shows the -log10(adj. p-values) of testing for associations between the factor scores and the condition or batch label. The upper panel shows the percentage of explained variance of each cell-type expression matrix recovered by the factor. c) Distribution of silhouette widths of each sample modeled by MOFA and scITD grouped by their condition or their technical label. Adjusted p-values < 0.05 of t-tests are shown. d) Loading scores of MOFA’s Factor 1 and scITD’s Factor 1 for POSTN and TTN across cell-types

**Supplementary Figure 2.**
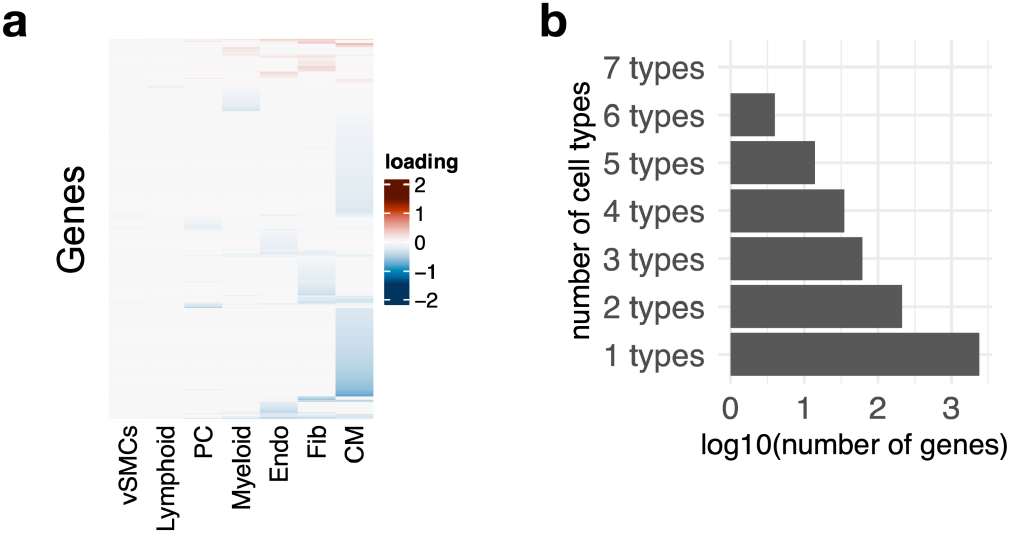
Multicellular program associated with myocardial remodeling. a) Gene weights of Factor 1 estimated from the human myocardial infarction single-cell data set across cell-types, representing the multicellular program associated with myocardial remodeling. b) Log10 number of genes belonging to distinct cell-type specific factor signatures

**Supplementary Figure 3.**
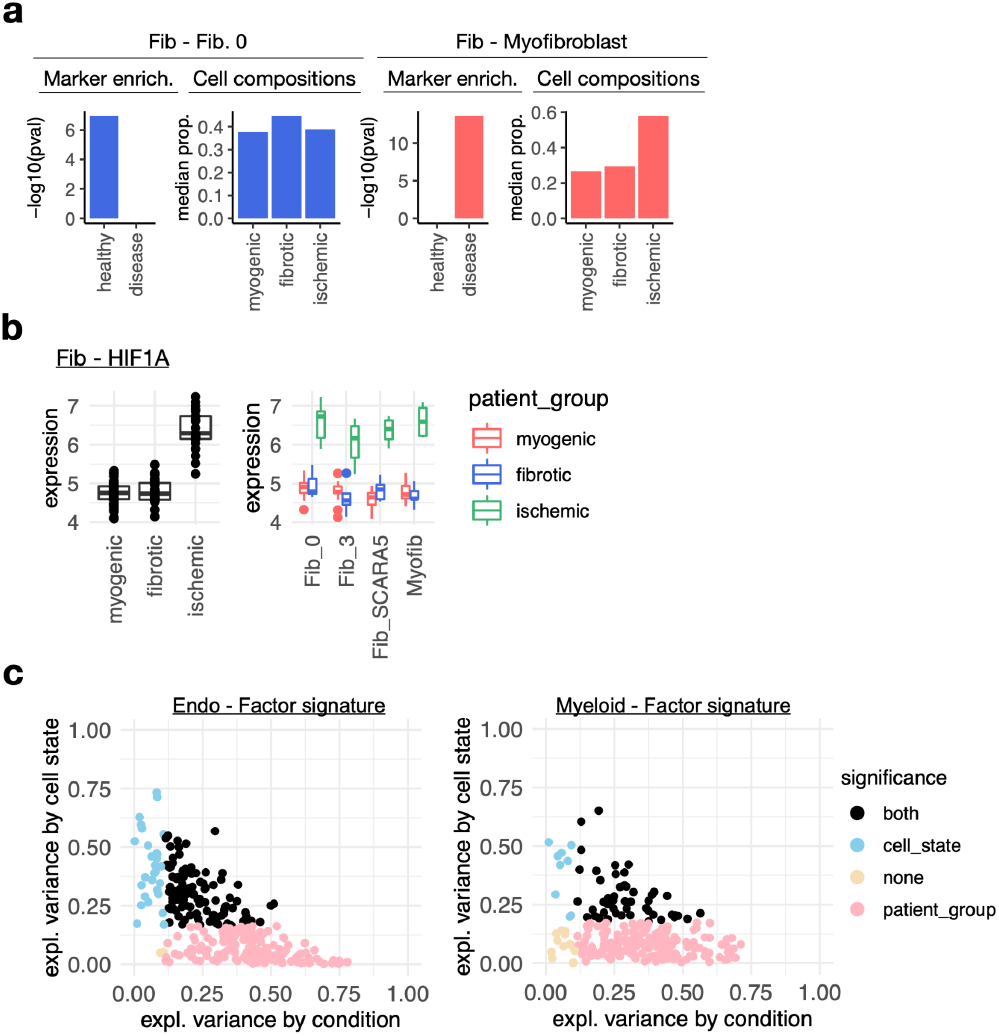
Multicellular program associated with myocardial remodeling. a) Cell-state markers captured in cell-type factor 1 signatures of Fibroblasts (Fib). In each panel is shown the adj. p-value of the enrichment of cell state markers of the indicated state in cell-type specific factor signatures (left) and the group median proportion of the cell-state across patient groups calculated from single-cell data (right). An example of the most overrepresented state in myogenic (left column) and ischemic (right column) samples is shown per cell-type. b) Distribution of log-normalized gene expression across samples of the human myocardial infarction atlas of HIF1A from pseudobulk profiles of Fibs. The left column is the pseudobulk expression of the gene for each sample, while the right column comes from the pseudobulk expression of the gene in a cell-state. c) Proportion of variance (eta-squared) of gene expression of genes belonging to the factor 1 signatures of endothelial (Endo) and myeloid cells explained by the patient’s conditions or cell states. Colors represent significance of the association of pseudobulk expression variability to patient’s conditions or cell states (ANOVA adj. p-value < 0.05).

**Supplementary Figure 4.**
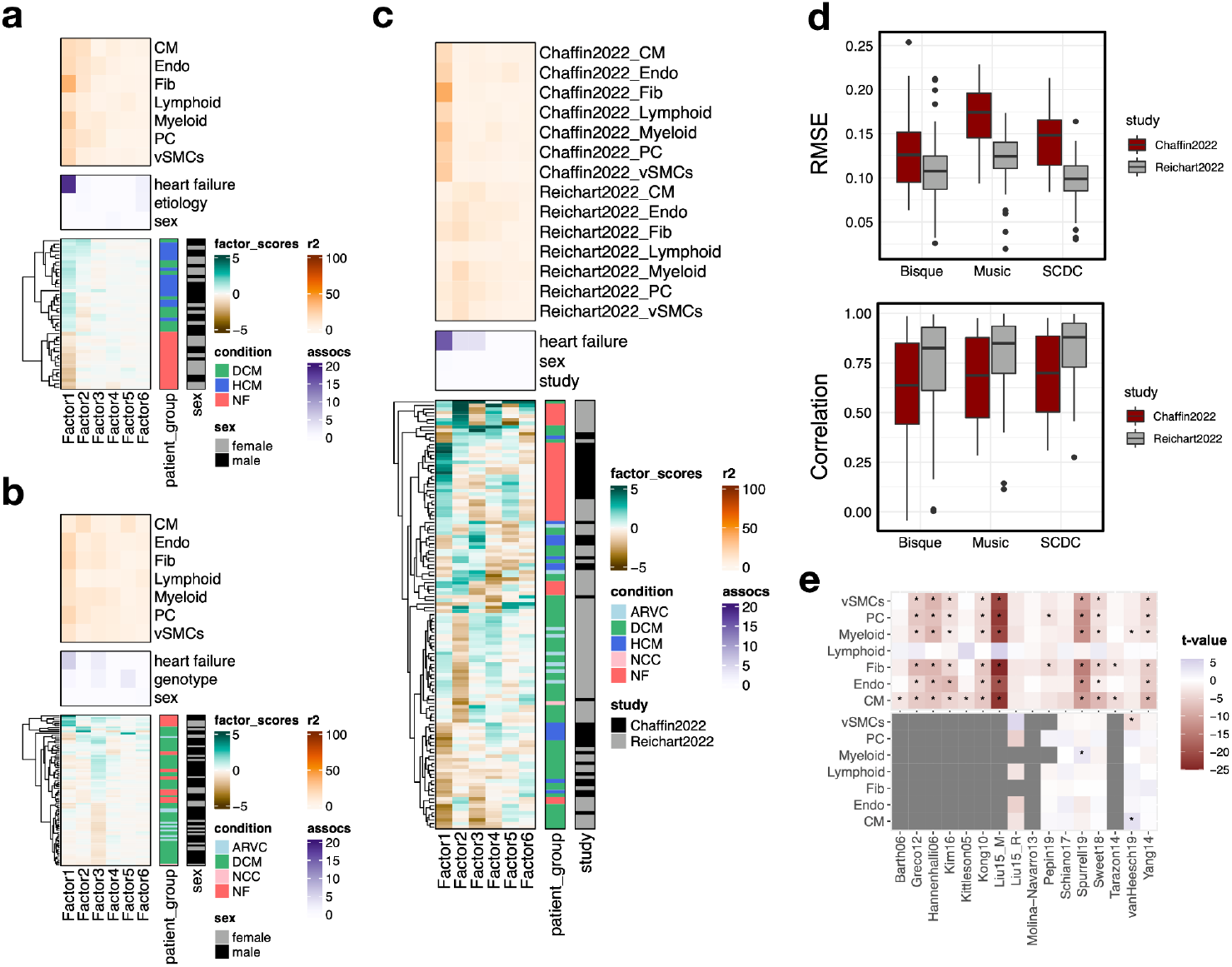
Multicellular factor analysis to integrate multiple single-cell cohorts and to deconvolute disease signals from bulk transcriptomics. a) Summary statistics of the multicellular factor analysis model for Chaffin2022. The lower panel shows the hierarchical clustering of the factor scores inferred by the MOFA model for the samples in the study. The patient’s condition and sex are indicated in each row. The middle panel shows the -log10(adj. p-values) of testing for associations between the factor scores and heart failure, etiology, and sex. The upper panel shows the percentage of explained variance of each cell-type expression matrix recovered by the factor. DCM = dilated cardiomyopathy, HCM = hypertrophic cardiomyopathy, NF = non-failing. b) Summary statistics of the multicellular factor analysis model for Reichart2022. The lower panel shows the hierarchical clustering of the factor scores inferred by the MOFA model for the samples in the study. The patient’s condition and sex are indicated in each row. The middle panel shows the -log10(adj. p-values) of testing for associations between the factor scores and heart failure, genotype, and sex. The upper panel shows the percentage of explained variance of each cell-type expression matrix recovered by the factor. ARVC = arrhythmogenic right ventricular cardiomyopathy, DCM = dilated cardiomyopathy, NCC = non-compaction cardiomyopathy, NF = non-failing. c) Summary statistics of the grouped multicellular factor analysis model of Chaffin2022 and Reichart2022. The lower panel shows the hierarchical clustering of the factor scores inferred by the MOFA model of the 121 samples. The patient’s condition, sex, and study labels are indicated in each row. The middle panel shows the -log10(adj. p-values) of testing for associations between the factor scores and heart failure, sex, or the study label. The upper panel shows the percentage of explained variance of each cell-type expression matrix recovered by the factor for each study separately. ARVC = arrhythmogenic right ventricular cardiomyopathy, DCM = dilated cardiomyopathy, HCM = hypertrophic cardiomyopathy, NCC = non-compaction cardiomyopathy, NF = non-failing. d) Evaluation of the performance of the cell-type deconvolution methods Bisque, Music and SCDC in the estimation of true cell compositions from pseudobulk samples of Chaffin2022 and Reichart2022. Upper panel shows the root-mean-squared-error and the lower panel the Pearson correlation to true values. e) T-values obtained from the comparison of heart failing vs non-failing samples across 16 independent bulk studies from ReHeaT using cell-type factor signatures (upper) or estimated cell-type compositions (lower). Stars highlight differences between conditions (adj. p-value <= 0.1). Gray tiles represent studies where the comparison was not possible (Microarray study or compositions all equal to 0). CM = cardiomyocytes, Fib = fibroblasts, Endo = Endothelial, vSMCs = vascular smooth muscle cells, PC = pericytes.

## Notes

https://zenodo.org/record/7660312#.Y_YxyuzMIeZ

https://github.com/saezlab/MOFAcell

### Competing Interest Statement

J.S.R. reports funding from GSK, Pfizer and Sanofi and fees from Travere Therapeutics, and Astex.

